# Multiplexed gene editing with a multi-intron containing *Cas9* gene in citrus

**DOI:** 10.1101/2023.12.15.571842

**Authors:** Poulami Sarkar, Jorge Santiago Vazquez, Mingxi Zhou, Amit Levy, Zhonglin Mou, Vladimir Orbović

## Abstract

The citrus industry holds significant economic importance in Florida, being one of the leading producers of oranges and grapefruits in the United States. However, several diseases, such as canker and huanglongbing along with natural disasters like hurricanes have rigorously affected citrus production, quality, and yield. Improving citrus through traditional breeding methods requires significant challenges due to time constraints and complexity in genetic enhancements. To overcome these limitations, several expression systems have been developed in clustered regularly interspaced short palindromic repeats (CRISPR)-associated protein 9 (CRISPR/Cas9) framework allowing for gene editing of disease-associated genes across diverse citrus varieties. In this study, we present a new approach employing a multi-intron containing *Cas9* gene plus multiple gRNAs separated with tRNA sequences to target the phytoene desaturase (*PDS*) gene in both ‘Carrizo’ citrange and ‘Duncan’ grapefruit. Notably, using this unified vector significantly boosted editing efficiency in both citrus varieties, showcasing mutations in all three designated targets. The implementation of this multiplex gene editing system with a multi-intron-containing *Cas9* plus a gRNA-tRNA array demonstrates a promising avenue for efficient citrus genome editing, equipping us with potent tools in the ongoing battle against HLB.

**Statements and Declarations:** *Competing interests:* The authors declare that they have no competing interests.

*Supplementary Information:* Supplementary File 1

## Introduction

Citrus stands as a vital fruit crop on a global scale and holds particular significance in Florida. The citrus industry in Florida has recently declined by 74 % and has resulted in California surpassing it as the foremost citrus-producing state (Weber et al. 2023; Bové 2006; Singerman and Rogers 2020; Haque et al. 2023). This weakening citrus production is attributed to various biotic stress, including bacterial and viral diseases such as Canker, Huanglongbing, and natural disasters like Hurricanes and freezes (Ferrarezi et al. 2020; Gottwald and Graham 2014; Gabriel et al. 2020; Harper and Cowell 2016). At present, numerous strategies are under development and being implemented to manage citrus diseases. Some of those include thermotherapy (Ghatrehsamani et al. 2019; Armstrong et al. 2021), trunk-injections with antimicrobials (Archer et al. 2023; Archer and Albrecht 2023), plant defense inducers (Hu et al. 2018), control of the insect vector (Snyder et al. 2022; Diniz et al. 2020), use of CUPS (citrus under protective screening) (Schumann et al. 2022) and eradication of HLB or canker symptomatic trees (Lee et al. 2015; Li et al. 2021; Bassanezi et al. 2009). However, these techniques are often ineffective, can pose risks to both food safety and the environment and may contribute to the emergence of antibiotic-resistant microorganisms (Shin et al. 2016; de Gracia Coquerel et al. 2023; Vashisth and Livingston 2020). Traditional breeding methods have proven effective in enhancing citrus cultivars but are limited to traits associated with fruit quality and constrained by time (Conti et al. 2021).

During the last two decades, genetic engineering methods have been successfully adopted to improve fruit quality, nutritional value, and disease tolerance. Genome editing via clustered regularly interspaced short palindromic repeats -associated protein 9 (CRISPR/Cas9) system is being widely applied to crops to induce disease/abiotic stress tolerance (Rai and Shekhawat 2014; Jia and Wang 2014; Jia et al. 2019; Li et al. 2019; Abdallah et al. 2022; Dutt et al. 2020). In assessing the effectiveness of the CRSPR/Cas9 technique, the phytoene desaturase (*PDS*) gene is a frequently targeted candidate (Qin et al. 2007). The disruption of this gene leads to a deficiency in carotenoid and chlorophyll production, resulting in a visibly distinct white albino phenotype that facilitates easy visualization and selection. There are several architectural parameters to design efficient *Cas9* gene and guide RNA sequences in a construct, and one of the most efficient is the combination of a multi-intron-containing *Cas9* gene plus multiplex gRNAs, which improves the efficiency as well as increases the number of targets (Grützner et al. 2021; Stuttmann et al. 2021).

This study introduces an efficient CRISPR/Cas9 vector with the multi-intron containing *Cas9* gene and three gRNAs separated by tRNA sequences, which facilitated successful editing of the *PDS* gene in two different citrus varieties, ‘Carrizo’ citrange (CAR) and ‘Duncan’ grapefruit (DUN). Our results demonstrated a high editing efficiency when coupled with the transformation process. This vector is a valuable tool for enhancing citrus genetic improvement and advancing precision breeding practices.

## Results and Discussion

The vector pAGM-PDS was made by inserting a gRNA-tRNA array containing three gRNAs targeting the citrus *PDS* gene into pAGM55273 (Fig. 1). The vector was transformed into *Agrobacterium* strain EHA105. This recombinant *Agrobacterium* was used to transform citrus epicotyl explants based on the previously published protocol (Orbović and Grosser 2015).

**Fig. 1.**
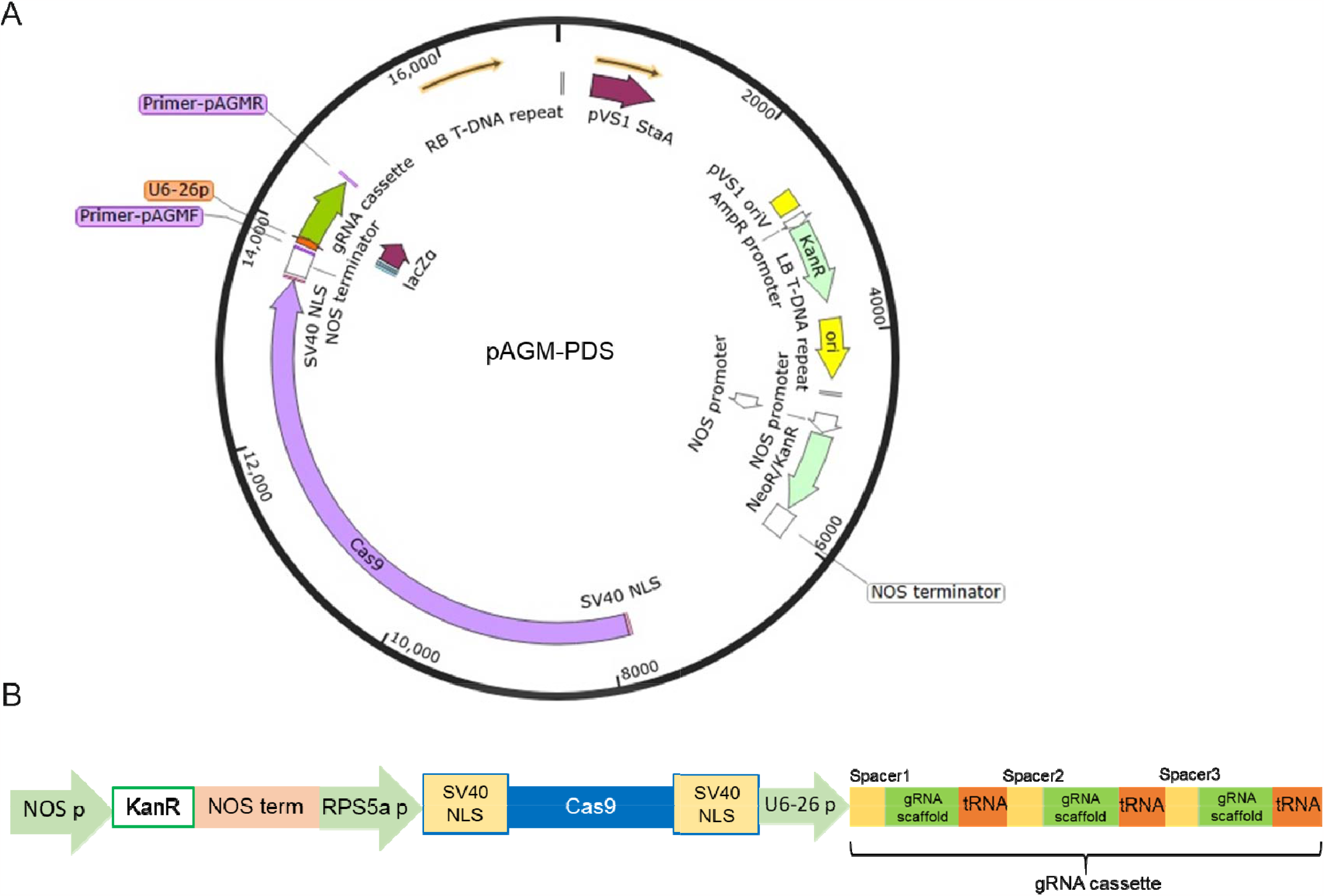
Construct pAGM-PDS used for editing with the gRNA-tRNA array into pAGM55273.

Due to high regeneration potential observed in CAR explants, their incubation in regeneration media (RM) without Kanamycin (–) resulted in uncontrolled shoot regeneration within a week. Under these conditions, no transgenic and/or edited shoots were observed. This outcome is most probably the result of faster and more vigorous growth of wild-type shoots that can impede the growth and survival of transgenic shoots. When CAR explants were incubated on kanamycin-containing media (+), they exhibited robust shoot regeneration with noticeable white shoot formation. A much lower shoot generation was recorded for DUN explants on Kanamycin-containing media during the initial month, followed by transfer to new RM plates with or without Kanamycin (Table 1). Consequently, we conducted experiments involving CAR by maintaining the explants in Kanamycin-containing media and DUN on RM media without kanamycin to have a balanced number of shoot regeneration with higher editing rate.

**Table 1:** Regeneration rates expressed as the number of shoots per explant for CAR and DUN explants incubated on RM media with (+) or without (-) Kanamycin for period of one month followed by additional two months. Numbers in parentheses are the total number of shoots/total

| Type of media | - → - | - → + | + → + | + → - |
| --- | --- | --- | --- | --- |
| **CAR** | 2.04 (249/122) | 1.18 (143/121) | 0.93 (111/120) | 1.3 (157/120) |
| **DUN** | 0.88 (106/120) | 0.61 (86/140) | 0.19 (23/120) | 0.14 (14/100) |

number of explants.

The initial appearance of the first white shoot in CAR was observed at 40 days post transformation (Fig. 2, upper panels), whereas in DUN, it was observed after 60 days (Fig. 3A). Our standard protocol for production of transgenic and/or edited shoots calls for inspection and counting of shoots, and cessation of experiments 35-40 days after co-incubation with *Agrobacterium*. However, in this study, the process extended beyond this timeframe and the development of the phenotypes resulting from the action of editing machinery within the shoot tissue took unexpectedly longer time (around 90 days). Also, some of the white CAR shoots developed from what were originally green shoots (Fig. 2). Promotion of development of white phenotype in CAR shoots due to the editing event was reported after the application of heat treatment to co-incubated citrus explants (LeBlanc et al, 2018). However, shoots presented in Figure 2 sprouted from explants that were not exposed to heat treatment. It was previously suggested that different promoters can affect the timing of expression of Cas9 nuclease and as a result frequency of production of chimeric shoots in citrus. A comparison was made between 35S, CmYLCV, and CsUbi promoters and the conclusion was that expression of *Cas9* under 35S control starts later and consequently more chimeric shoots are produced (Huang et al. 2022). In the pAGM-PDS vector, *Cas9* is under control of *RPS5a* promoter that may also induce *Cas9* expression once the shoots are already formed. Both of these phenomena; extended time for development of phenotype and non-promoted development of phenotype may be the result of the employment of *RPS5a* promoter and deserve further exploration.

**Fig. 2.**
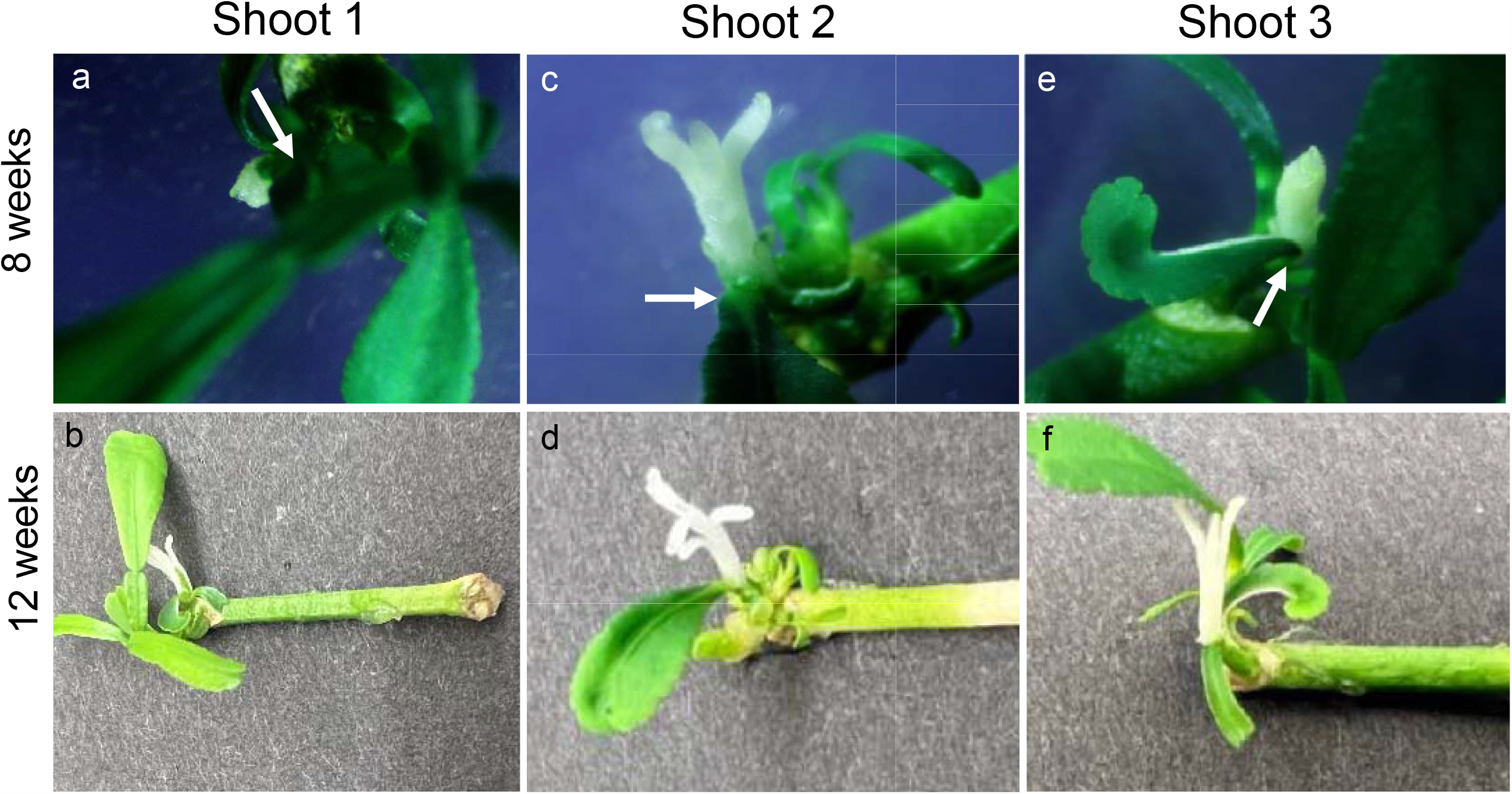
Phenotype development in CAR edited shoots. Photographs on the upper panels depict very young white shoots appearing from originally green shoots (labeled with white arrows) observed around 9-week post transformation (a, c, e). Photographs on the lower panels show well developed, fully white shoots observed around 12-week post transformation (b, d, f).

**Fig. 3.**
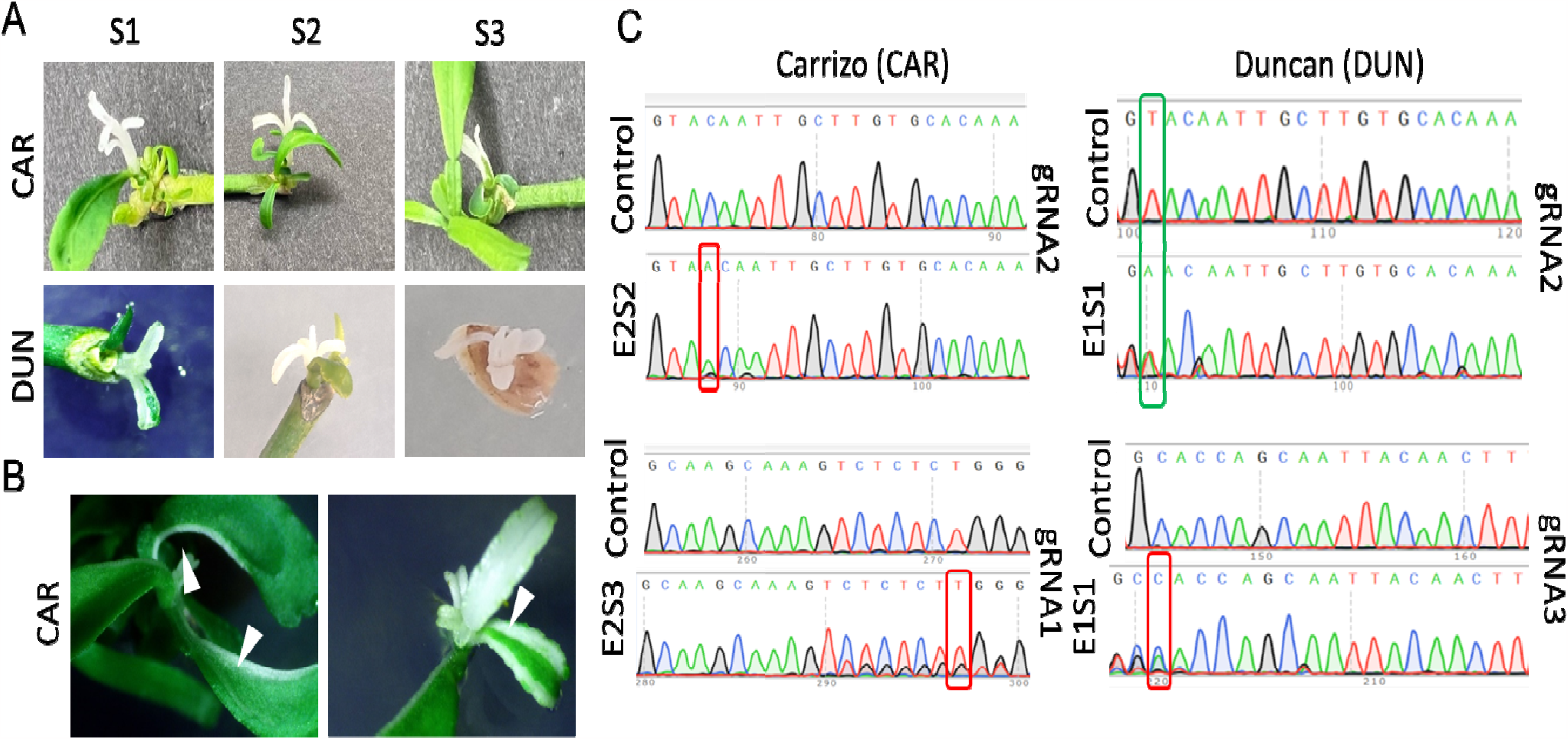
Detection of editing in the white shoots. A) White shoots as regenerated from explants incubated on Regeneration media with Kan for CAR and without Kanamycin for DUN (S1-3 denotes shoot numbers). B) Chimeric shoots as observed on CAR shoots regenerated from explants transformed with *Agrobacterium* EHA105-pAGM-PDS. C) Representative mutations in the target gRNA sequences of the white shoots. (E: Experiment number and S: White Shoot number.)

The distinct white shoots were allowed to grow for an additional two week before further analyses. Several chimeric shoots observed only in CAR are shown in Fig. 3B. Subsequently, a pair of *PDS*-specific primers (Table 5) were used to amplify the spacer sequence regions from genomic DNA isolated from the white shoots and the resulting PCR products were sequenced. All the shoots with edited *PDS* sequence were stably transformed, as confirmed by PCR analysis with the primers pAGMF-pAGMR (Fig. 4).

**Fig. 4.**
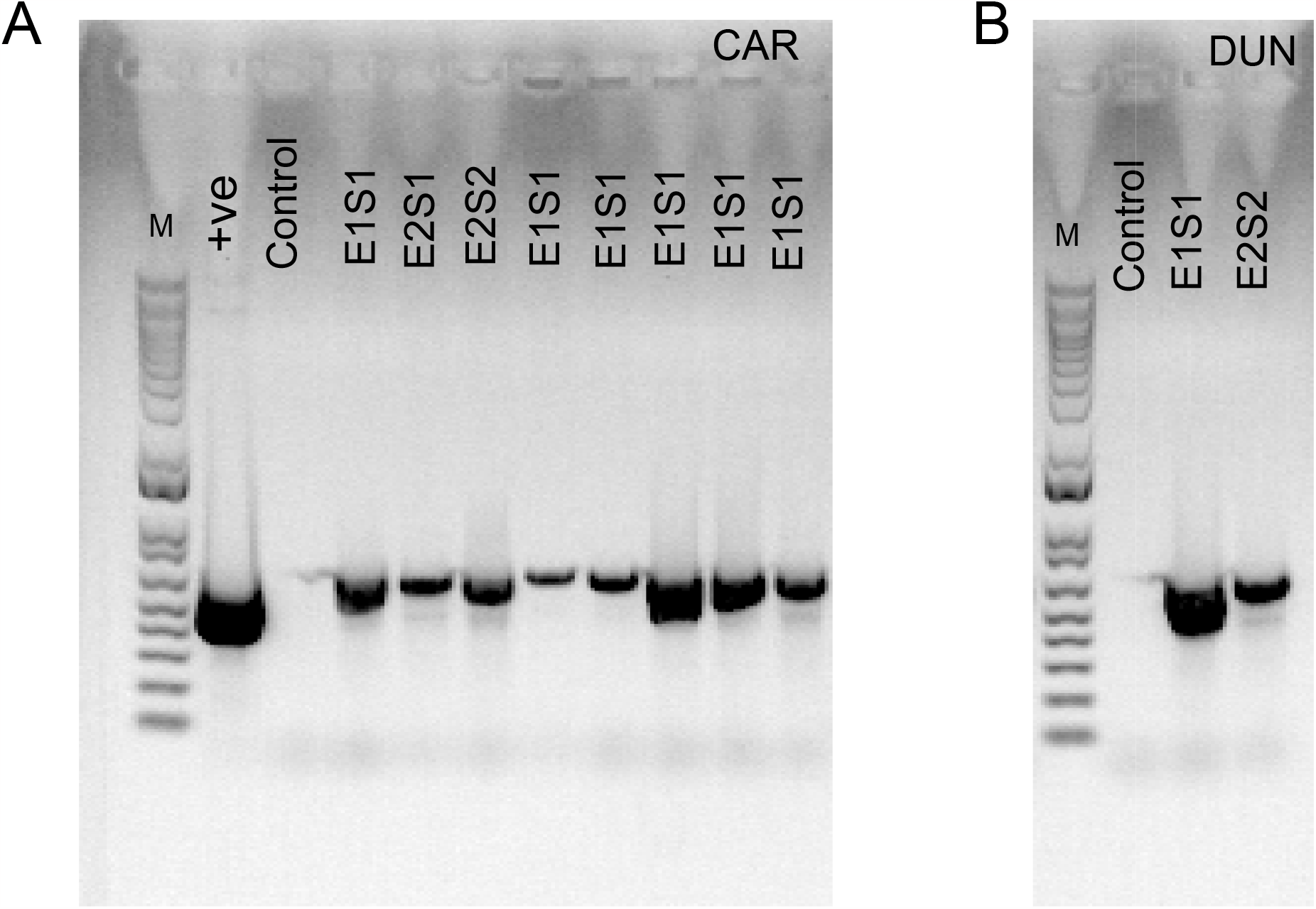
PCR analysis of all the white shoots compared with the control shoots generated from both A) CAR and B) DUN explants. (E: Experiment number and S: White Shoot number, M: 100bp Plus DNA ladder)

The chimeric shoots were excluded from the sequencing analysis as they could yield genomic DNA with non-edited sequences. The editing efficiency in CAR averaged to around 3.04 ± 1.37 % (4.5% in one experiment) with mutations observed in all three gRNA targets. This efficiency is high considering that the transformation efficiency in CAR is often below 10% (Dominguez et al. 2022; Ballester et al. 2008; Dutt and Grosser 2009). In DUN, the editing efficiency was around 0.54 ± 0.39 %, which is lower than that in CAR (Table 2). Most of the white shoots had a point mutation, specifically insertion, although substitutions were also found in some targets (Fig. 2C and Table 3). The multi-intron-containing *Cas9* gene has been shown to be highly efficient in Arabidopsis and tobacco (Stuttmann et al. 2021; Schindele et al. 2022; Grützner et al. 2021) and the gRNA-tRNA array with multiple gRNAs to target several genomic sites has been shown to elevate the expression levels of multiplexed gRNAs within the same transcript (Jiang et al. 2023; Zhang et al. 2019; Xie et al. 2015). The editing rate observed in this study, coupled with the ability of this system to successfully edit multiple targets in two distinct citrus varieties, holds promise for manipulation of multiple genes associated with HLB symptoms.

**Table 2:** Editing rate in Carrizo (on Kan) and Duncan grapefruit (without Kan) with pAGM-PDS construct.

| Experiment<br>(E) | #White<br>shoots (S) | #Chimeric<br>Shoots | #Total shoots/ Total<br>Explant | Editing rate |
| --- | --- | --- | --- | --- |
| **CAR-E1** | 1 | 0 | 82/100 | 1.2 % |
| **CAR-E2** | 4 | 2 | 176/120 | 3.41 % |
| **CAR-E3** | 3 | 4 | 153/120 | 4.5 % |
| **DUN-E1** | 1 | 0 | 106/120 | 0.94 % |
| **DUN-E2** | 2 | 0 | 296/150 | 0.67 % |
| **DUN-E3** | 0 | 0 | 154/120 | 0 % |

**Table 3:**
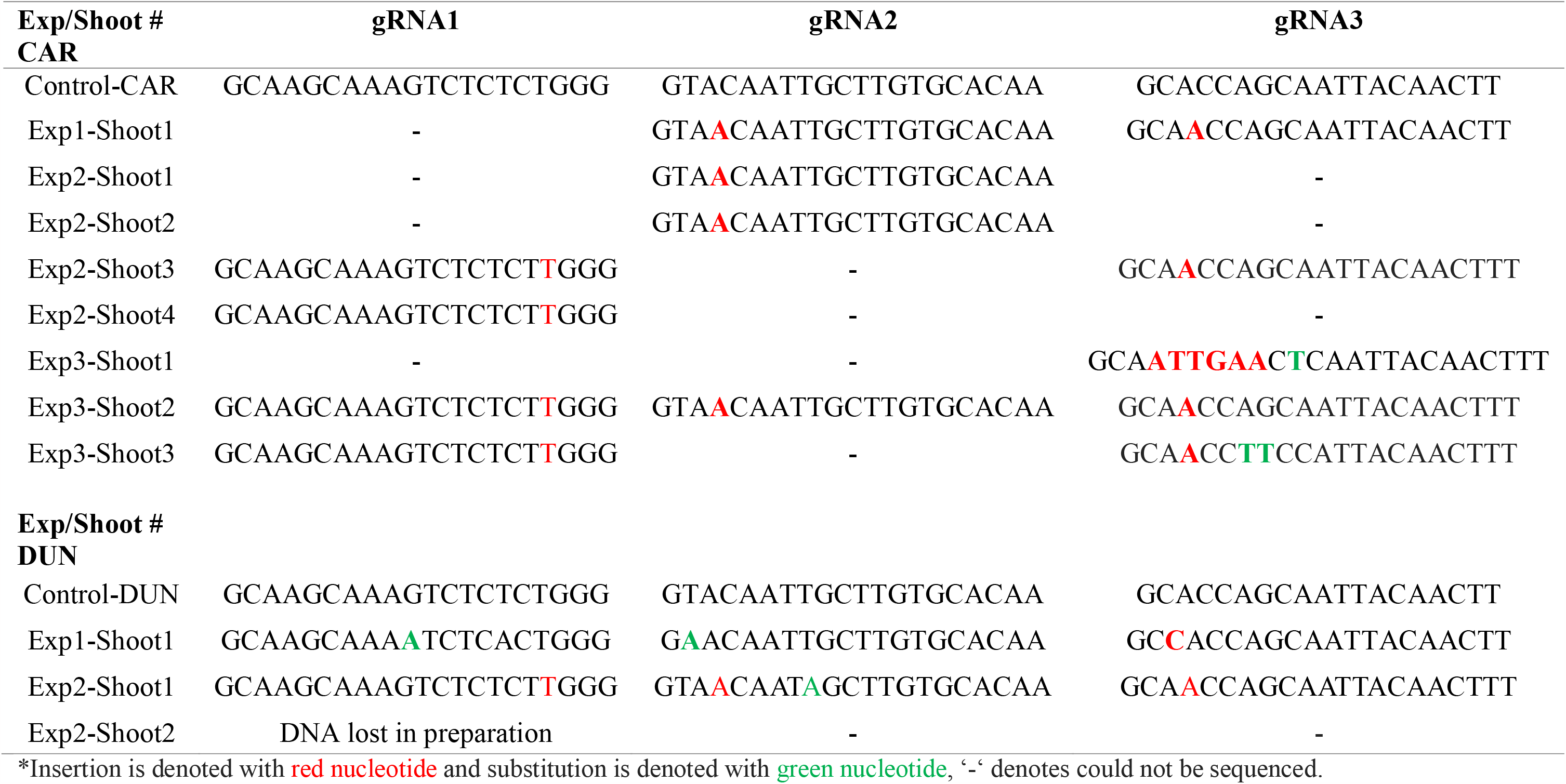
Sequences of target gRNA after editing with pAGM-PDS.

To increase the genome editing rate, CAR explants were subjected to heat treatment as previously reported for citrus (LeBlanc et al. 2018). Although this treatment increased efficiency of editing in two other crops as well (Blomme et al. 2022; Milner et al. 2020), no significant change was recorded in our experiments (Table 4).

**Table 4:**
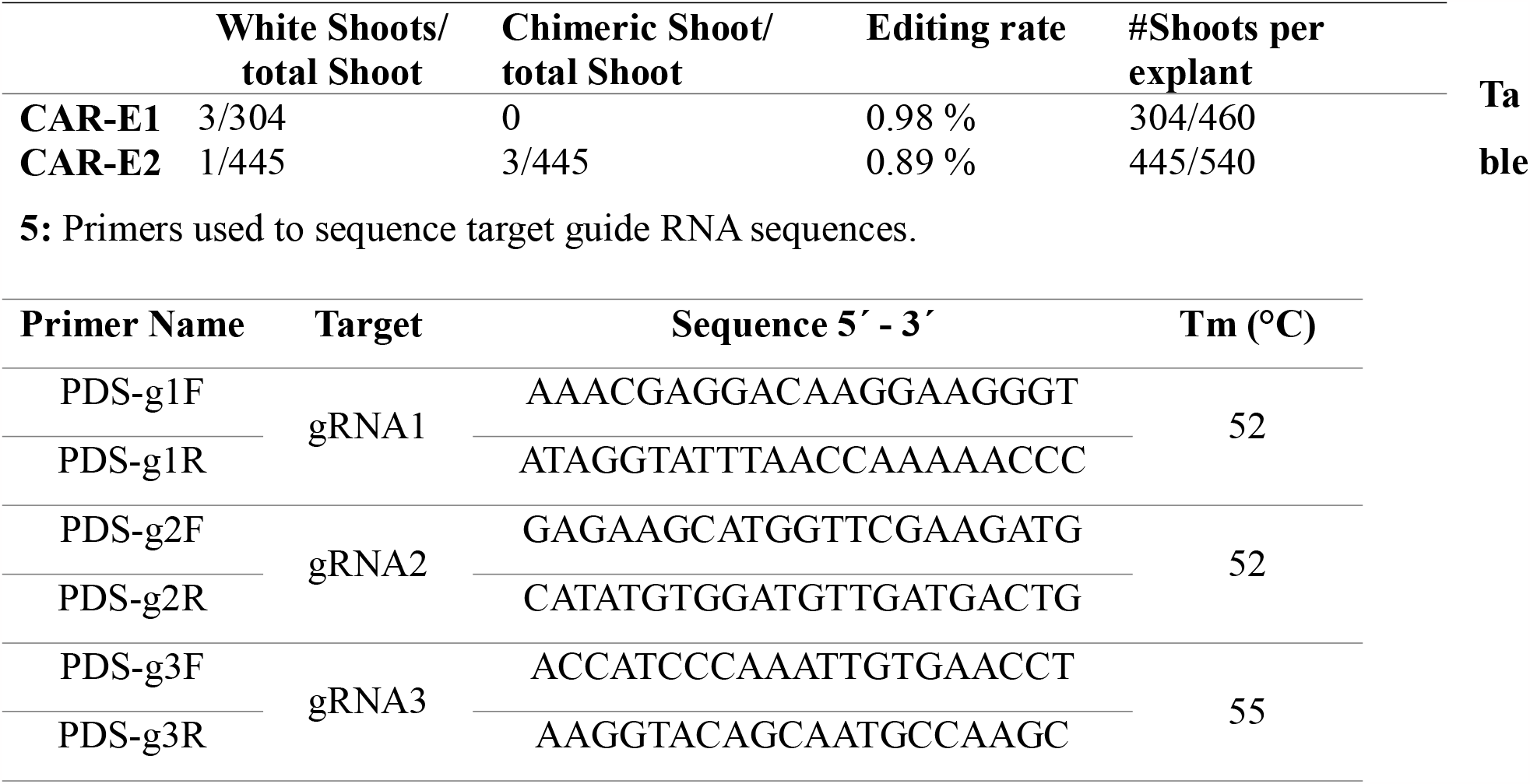
Editing rate and regeneration rate with heat treatment in CAR maintained on Kanamycin containing RM media.

In summary, we developed an efficient vector capable of targeting three different targets either within the same gene or in different genes. Although CAR is reported to have a high transformation efficiency, our study found a high editing efficiency up to 4.5 % in conjunction with the transformation process. This high efficiency is attributed to the utilization of a multi-intron-containing *Cas9* and a gRNA-tRNA array. Further refinement of this vector is required for effective generation of multiple genome-edited non-transgenic citrus trees. This advancement holds great potential in contributing to the improvement of citrus varieties to combat HLB.

## Materials and methods

### Constructs used

To prepare the construct pAGM-PDS, we used pAGM55273 (Addgene plasmid #153211) as the backbone that contains the *Cas9* gene with 13 introns and kanamycin selection cassette for plant transformation. We introduced a gRNA-tRNA array with three gRNAs targeting the *Citrus sinensis PDS* gene (>spacer1: TTGTGCACAAGCAATTGTAC, >spacer2: GCAAGCAAAGTCTCTCTGGG and >spacer3: AAAGTTGTAATTGCTGGTGC). A gRNA scaffold-tRNA template was synthesized by GenScript and cloned into pBlueScriptII (SK) with SacI and SalI (Fig. 1A, Supplementary material 1). The gRNA-tRNA array was introduced into pAGM55273 using Golden Gate Assembly (BsaI) with the primers listed in the Supplementary material 1.

### Agrobacterium strains and plant material

The construct pAGM-PDS was transformed into *Agrobacterium tumefaciens* strain EHA105 and this transformed bacterium was used for the plant transformation experiments on ‘Carrizo’ citrange [*Poncirus trifoliata* (L.) Raf. ×◻*Citrus sinensis* (L.) Osbeck] (CAR) and ‘Duncan’ grapefruit (Citrus paradisi Macf.) (DUN). The primers used to confirm the vector are as follows: pAGMF: GACGCTACTAGAATTCGAGC, and pAGMR: TCCGCGACATTCTAGAACTC.

### Plant transformation

CAR and DUN seeds were peeled, surface-sterilized in 30 % bleach for 12 minutes followed by rinsing with water, thrice. The sterilized seeds were then planted in tubes containing MS medium (Murashige and Skoog 1962). Thirty days after germination, the seedlings were transferred to light to etiolate for 3 days before being cut for explants. 2cm stem cuts from these seedlings were used as explants and were placed in liquid co-cultivation medium (CCM) (Orbović and Grosser 2007) for 2-3 h prior to transformation with *Agrobacterium* EHA105-pAGM-PDS. This was followed by soaking the explants in recombinant *Agrobacterium* containing the construct (EHA105-pAGM-PDS) (OD= 0.7) for 3min, then was incubated on plates with CCM-agar medium for 2 days in dark. The explants were transferred to regeneration medium (RM) (Orbović and Grosser 2007) with antibiotic Cefotaxime (330 mg/L) to eliminate *Agrobacterium* and with or without Kanamycin (Kan). Control shoots were regenerated from non-transformed explants on RM medium. The explants were incubated in dark for 1 week before moving to growth chamber to regenerate for 4 months and were observed every week for white shoot development.

For heat treatment, the RM plates containing the *Agrobacterium*-transformed explants were incubated at 35 °C for 8h for three consecutive days, followed by incubation at 24 °C in a growth chamber.

### DNA isolation and PCR analysis of mutations

The generated white shoots from each experiment were used for DNA isolation followed by target gRNA Sanger’s sequencing. Briefly, the white shoots were crushed in liquid nitrogen and were used for DNA isolation using CTAB (Doyle et al. 1987). Non-transformed control shoots were used as a control to compare the mutations. PCR was performed using Phusion high-fidelity DNA polymerase (Thermo Fisher Scientific) and three specific primers for each gRNA sequence (Table 5).

## Acknowledgements

This research was supported by the USDA/NIFA Emergency Citrus Disease Research and Extension Program.

## Notes

### Competing Interest Statement

The authors have declared no competing interest.

