## Supplementary material for "Multiplexed gene editing with a multi-intron containing *Cas9* gene in citrus"

**Construction of pAGM-PDS**

For CRISPR/Cas9 vector pAGM55273, using the pMOD_B_M vector as template.

For citrus PDS gene spacers:

>gRNA1

GCAAGCAAAGTCTCTCTGGG (909-928)

>gRNA2

TTGTGCACAAGCAATTGTAC (18349)

>gRNA3

AAAGTTGTAATTGCTGGTGC (258)

Primers:

F1:

5’ATATATGGTCTC**G**ATT**GCAAGCAAAGTCTCTCTGGG**GTTTAAGAGCTATGCTGGAAACAG3’

R1:

5’ATTATTGGTCTC**GGCTTGTGCACAA**TGCACCAGCCGGGAATCG3’

F2:

5’ATATATGGTCTC**GAAGCAATTGTAC**GTTTAAGAGCTATGCTGGAAACAG3’

R2:

5’ATTATTGGTCTC**GAAACGCACCAGCAATTACAACTTT**TGCACCAGCCGGGAATCG3’

Sequencing result:

NNNNNNNNNTGANCATCTTCAAAGTCCCACATCGCTTAGATAAGAAAACGAAGCTGAGTTTATATACAGCTAGAGTCGAAGTAGTGATTGCAAGCAAAGTCTCTCTGGGGTTTAAGAGCTATGCTGGAAACAGCATAGCAAGTTTAAATAAGGCTAGTCCGTTATCAACTTGAAAAAGTGGCACCGAGTCGGTGCAACAAAGCACCAGTGGTCTAGTGGTAGAATAGTACCCTGCCACGGTACAGACCCGGGTTCGATTCCCGGCTGGTGCATTGTGCACAAGCAATTGTACGTTTAAGAGCTATGCTGGAAACAGCATAGCAAGTTTAAATAAGGCTAGTCCGTTATCAACTTGAAAAAGTGGCACCGAGTCGGTGCAACAAAGCACCAGTGGTCTAGTGGTAGAATAGTACCCTGCCACGGTACAGACCCGGGTTCGATTCCCGGCTGGTGCAAAAGTTGTAATTGCTGGTGCGTTTTAGAGCTAGAAATAGCAAGTTAAAATAAGGCTAGTCCGTTATCAACTTGAAAAAGTGGCACCGAGTCGGTGCTTTTTTTTCGCTTTACGAATTCCCATGGGGAGTTCTAGAATGTCGCGGAACAAATTTTAAAACTAAATCCTAAATTTTTCTAATTTTGTTGCCAATAGTGGATATGTGGGCCGTATAGAAGGAATCTATTGAAGGCCCAAACCCATACTGACGAGCCCAAAGGTTCGTTTTGCGTTTTATGTTTCGGTTCGATGCCAACGCCACATTCTGAGCTAGGCAAAAAACAAACGTGTCTTTGAATAGACTCCTCTCGTTAACACATGCAGCGGCTGCATGGTGACGCCATTAACACGTGGCCTACAATTGCATGATGTCTCCATTGACACGTGACTTCTCGTCTCCTTTCTTAATATATCTAACAAACACTCCTACCTCTTCCAAAATATATACACATCTTTTTGATCAATCTCTCATTCAAATCTCATTCTCTCTANTAAACAANAACAAAAAAATGGCGGATACAGCTANAGGAACCCATCACGATATCATCGGNANAANATCAGTACCCGATGATGGGNCCNNNN

Primers for colony PCR: vector pAGM55273

pAGMF: GACGCTACTAGAATTCGAGC

pAGMR: TCCGCGACATTCTAGAACTC

Empty vector: 924

3 gRNAs: 717

The pMOD_B_M vector:

The following Sequence in pBluescript II SK(+) cloned with SacI and SalI

**GAGCTC**CACCTGCCAGGGGACCGACTTGCCTTCCGCACAATACATCATTTCTTCTTAGCTTTTTTTCTTCTTCTTCGTTCATACAGTTTTTTTTTGTTTATCAGCTTACATTTTCTTGAACCGTAGCTTTCGTTTTCTTCTTTTTAACTTTCCATTCGGAGTTTTTGTATCTTGTTTCATAGTTTGTCCCAGGATTAGAATGATTAGGCATCGAACCTTCAAGAATTTGATTGAATAAAACATCTTCATTCTTAAGATATGAAGATAATCTTCAAAAGGCCCCTGGGAATCTGAAAGAAGAGAAGCAGGCCCATTTATATGGGAAAGAACAATAGTATTTCTTATATAGGCCCATTTAAGTTGAAAACAATCTTCAAAAGTCCCACATCGCTTAGATAAGAAAACGAAGCTGAGTTTATATACAGCTAGAGTCGAAGTAGTGATTGGCAACAAAGCACCAGTGGTCTAGTGGTAGAATAGTACCCTGCCACGGTACAGACCCGGGTTCGATTCCCGGCTGGTGCAG**GAGACG**GTTTAAGAGCTATGCTGGAAACAGCATAGCAAGTTTAAATAAGGCTAGTCCGTTATCAACTTGAAAAAGTGGCACCGAGTCGGTGCAACAAAGCACCAGTGGTCTAGTGGTAGAATAGTACCCTGCCACGGTACAGACCCGGGTTCGATTCCCGGCTGGTGCA**CGTCTC**GGTTTAAGAGCTATGCTGGAAACAGCATAGCAAGTTTAAATAAGGCTAGTCCGTTATCAACTTGAAAAAGTGGCACCGAGTCGGTGCTTTTTTTTGCAAAATTTTCCAGATCGATTTCTTCTTCCTCTGTTCTTCGGCGTTCAATTTCTGGGGTTTTCTCTTCGTTTTCTGTAACTGAAACCTAAAATTTGACCTAAAAAAAATCTCAAATAATATGATTCAGTGGTTTTGTACTTTTCAGTTAGTTGAGTTTTGCAGTTCCGATGAGATAAACCAATACCGGTGACGCAGGTG**GTCGAC**

AACCT: U6-26p

AACAAAGC: tRNA

GTTTAAGAGC: relaxed gRNA scaffold

CGTCTC: Esp3I
